# Sensory discrimination of chemical and temperature stimuli in the acoel *Symsagittifera roscoffensis*

**DOI:** 10.1101/2025.01.28.635210

**Authors:** Nikita Komarov, Christopher Aeschbacher, Laurent Sauterel, Evan Zuercher, Xavier Bailly, Pedro Martinez, Simon G. Sprecher

## Abstract

Environmental cues provide critical sensory information for the survival of animals. Understanding how distinct sensory cues elicit or modulate certain behaviour thus provides insights into the neuronal adaptations to rapid and continuous changes in the surrounding world. Intertidal ecosystems are a particularly exposed to environmental fluctuations. Due to changing exposure to seawater, animals are subjected to continuous fluctuations of temperature or salinity during the course of day-night and tidal cycles. Animals in the intertidal environment show physiological and behavioural adaptations to these changes. When the acoel *Symsagittifera roscoffensis* is exposed to daylight during the tidal cycle, these animals are found at the surface of sandy beaches, which enhance the exposure of their photosynthetic algal (*Tetraselmis convolutae*) symbionts to light. Moreover, *S. roscoffensis* shows a strong positive phototactic displacement as well as both positive and negative geotaxis, both being evolved behavioural adaptations to enhance light-exposure for its photosymbiont. Currently little is known about other sensory systems and their functions in *S. roscoffensis*. In this study, we probe sensory capabilities of *S. roscoffensis* focusing on chemical and temperature cues. Our findings support that *S. roscoffensis* shows avoidance behaviours to increased temperature and high salinity, preferring cooler environments with lower salinity.

## Introduction

While many animals utilise external sensory organs to determine the nature of their surroundings, the compositional, morphological and anatomical organization of sensory organs varies greatly in different animal clades. Sensory systems evolved to transduce signals of their surroundings into neuronally encoded information, which are subsequently integrated and processed in the central nervous system to ultimately elicit or modulate the behaviour of an animal.

The highly variable environmental physical and chemical conditions of intertidal ecosystems thus likely provide crucial sensory cues for animals. While desiccation as well as fluctuations in temperature and salinity are stressors for animals living in the intertidal zones, the direct exposure to light also provides advantages for certain physiological adaptations. One example of such specialisation can be found in the acoel worm *Symsagittifera roscoffensis*. This millimetre-scale worm lives in the intertidal zones of the Atlantic coasts of Europe, in a photosymbiotic relationship with the algae species *Tetraselmis convolutae* (Parke & Manton), and has captivated marine research since the 19^th^ century (Geddes, 1879; Keeble, 1911). Phylogenetically, it belongs to the phylum Xenacoelomorpha (Philippe et al., 2011) that diversified very early on within the Bilaterian lineage (Achatz et al, 2013(Achatz et al., 2013), though the precise phylogenetic placement of Xenacoelomorpha remains debated. As one of the few well-known bilaterian organisms capable of photosynthesis through photosymbiosis, *S. roscoffensis* challenges some conventional metabolic principles, born from the need to integrate processes occurring in two organisms under changing (daily or seasonally) environments, all of which has resulted in significant attention garnered from the fields of marine biology, ecology, and physiology (Arboleda et al., 2018; Bailly et al., 2021). However, despite receiving a range of nutrients and energy from its symbiont (Selosse, 2000), little is known about whether *S. roscoffensis* requires additional external nutrients (possible mixotrophy) or cues to thrive in this changing environment; and if so, how a hypothetical chemosensory system is used to forage for them. While lacking obvious or canonical chemosensory organs (only candidates: (Bery et al., 2010; Sakagami et al., 2024; Todt and Tyler, 2007)), *S. roscoffensis* possesses light-sensitive rhabdomeric eyes and a gravity-sensing statocyst (Bailly et al., 2014; Lanfranchi, 1990; Sprecher et al., 2015; Yamasu, 1991). While the strong phototactic behaviour as well as geotaxis of *S. roscoffensis* has been well investigated (Sprecher et al., 2015; Thomas et al., 2024), comparatively little is known about other sensory capacities.

Here, we try to remedy this lack of information by investigating the chemosensory and thermosensory sensitivities of *S. roscoffensis* by examining a variety or chemical and environmental modalities to determine the functional significance of identified sensory sensilla (Bery et al., 2010; Martínez et al., 2017).

Given that this worm lives and aquatic, intertidal, environment, where changes of physical parameters or chemical composition are most probable (e.g. light intensity, UV radiation, temperature, oxygen, and salinity (Adey and Loveland, 2007), we hypothesised that *S. roscoffensis* should be capable of sensing not only environmental stimuli such as temperature, and salinity, but are also able to respond to chemical cues, including those emitted by their internal symbionts (and their free-living relatives) or other species occupying their same ecological niche.

## Results

The chemosensory behaviors in *S. roscoffensis* remains currently understudied. In order to gain insights into the chemosensory capacity of *S. roscoffensis* we first performed two-choice assays to assess navigational responses when exposed to putatively relevant chemical cues. As first set of experiments we investigated the chemosensory behavioural response to a set of amino acids (AA) known to be used as neurotransmitter-precursors or neurotransmitters in both invertebrates and vertebrates (Roshchina, 2016). While the dependence of *S. roscoffensis* on its symbiont to provide essential neurotransmitters remains unknown, it has been previously shown that glutamate and aspartate, two amino acids that may act as neurotransmitters (Dingledine and McBain, 1999), were only detected in *S. roscoffensis* with symbionts, while it was absent in juvenile worms that did not yet harbour its symbiontic algae (Boyle and Smith, 1975). Other neurotransmitter AAs such as glycine, or neurotransmitter precursor AAs such as tyrosine (dopamine and noradrenaline precursors), tryptophan (serotonin precursor), were not tested for in previous studies. Given the conserved nature of these neurotransmitter systems in metazoans (Roshchina, 2016), it can be presumed that they are also present in Acoels. Hereby, we tested whether *S. roscoffensis* is able to sense and respond to the presence of these AA (glutamate, aspartate, tryptophane, tyrosine and glycine) in the environment (water column) using a two-choice navigation assay, measuring preference indices at 2-, 5-, and 10-minute timepoints. We placed animals on dishes containing agarose (control), or dishes containing half agarose and half agarose supplemented with the given amino acid (figure 1B). During 10-minute trials, we observed that the preference indices for glutamine (PI=-0.001, p=0.9996) and tyrosine (PI=-0.1, p=0.74) largely match those of the plain agarose control, where animals did not prefer either side (PI=0.03). For the amino acid aspartate, we observed a mild preference, albeit not showing statistically significant differences from the control (10min, PI=0.2, p=0.36, figure 1B, C). For glycine, we observed that there was a moderate attraction across the longer timescale (10min, PI=0.15, p=0.86, figure 1C). Tryptophan, however, evoked the strongest response, with significance being reached above the 5-minute timepoint (PI=0.46, p<0.0001 figure 1B, C).

**Figure 1:**
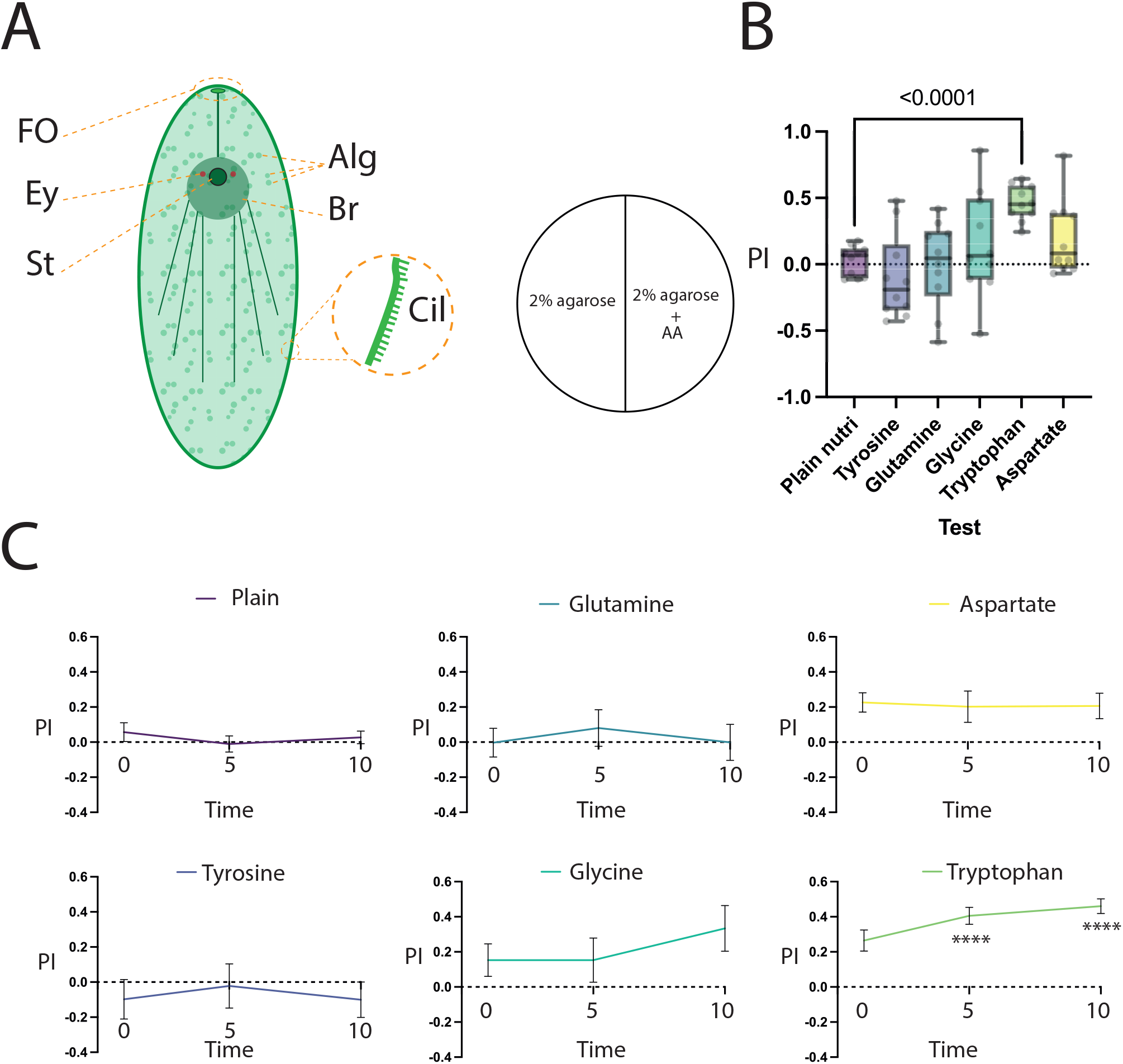
chemosensory preference of *S. roscoffensis* to neurotransmitter-related amino acids. A: Overview of *Symsagittifera roscoffensis* outlining the symbiont algae (Alg), brain (Br), eye spots (Ey), the gravisensory statocyst (St) and the putatively chemosensory frontal organ (FO). B: left: diagram of the behaviour arena indicating a two-choice set-up with one half containing plain agarose, and the other half containing agarose with dissolved test substance, in this case the indicated amino acid (AA). Right: preference indices for indicated test amino acids compared with the “nutrient plain” agarose control (purple) at the 5-minute timepoint, all trials displayed with min-max bars. C: Summary of the PIs shown as time-series across tested AAs, mean with SEM. N=10 for all tests. **** - p<0.0001, ns where not shown, Brown-Forsythe and Welch ANOVA test with Dunnet’s T3 multiple comparisons test against the control group.

Considering the dynamic nature with varying salinity in intertidal waters due to evaporation and temperature changes during low tides, it has been shown that during low tides the salinity of sea water can change by up to 300% (Geng et al., 2016), we then asked whether changes in salinity of the substrate would result in a preference phenotype.

Here, the animals were given a choice between a plain agarose/dH_2_O substrate and agarose with dissolved artificial sea water (ASW) adjusted to 34ppm salinity using a refractometer, to reflect the salinity of the ambient sea water measured outside the Roscoff marine station (figure 2A). We observed one-fold ASW (34ppm) was mildly, albeit insignificantly, aversive (10 min, PI=-0.22, p=0.09) compared with the environmental sea water control, and across all time points (figure 2B). However, we observed that two-fold ASW concentration (68ppm) was significantly aversive across the time series, reaching significance after the 5-minute timepoint (mean preference index -0.45, p<0.0001 vs control) (figure 2B). The concentration-dependent responses to variations in salinity thus allow us to assume a sensory ability to perceive high salt concentrations. This potentially reflects an ability to adapt to the dynamic variations in the intertidal environments where *S. roscoffensis* is found.

**Figure 2:**
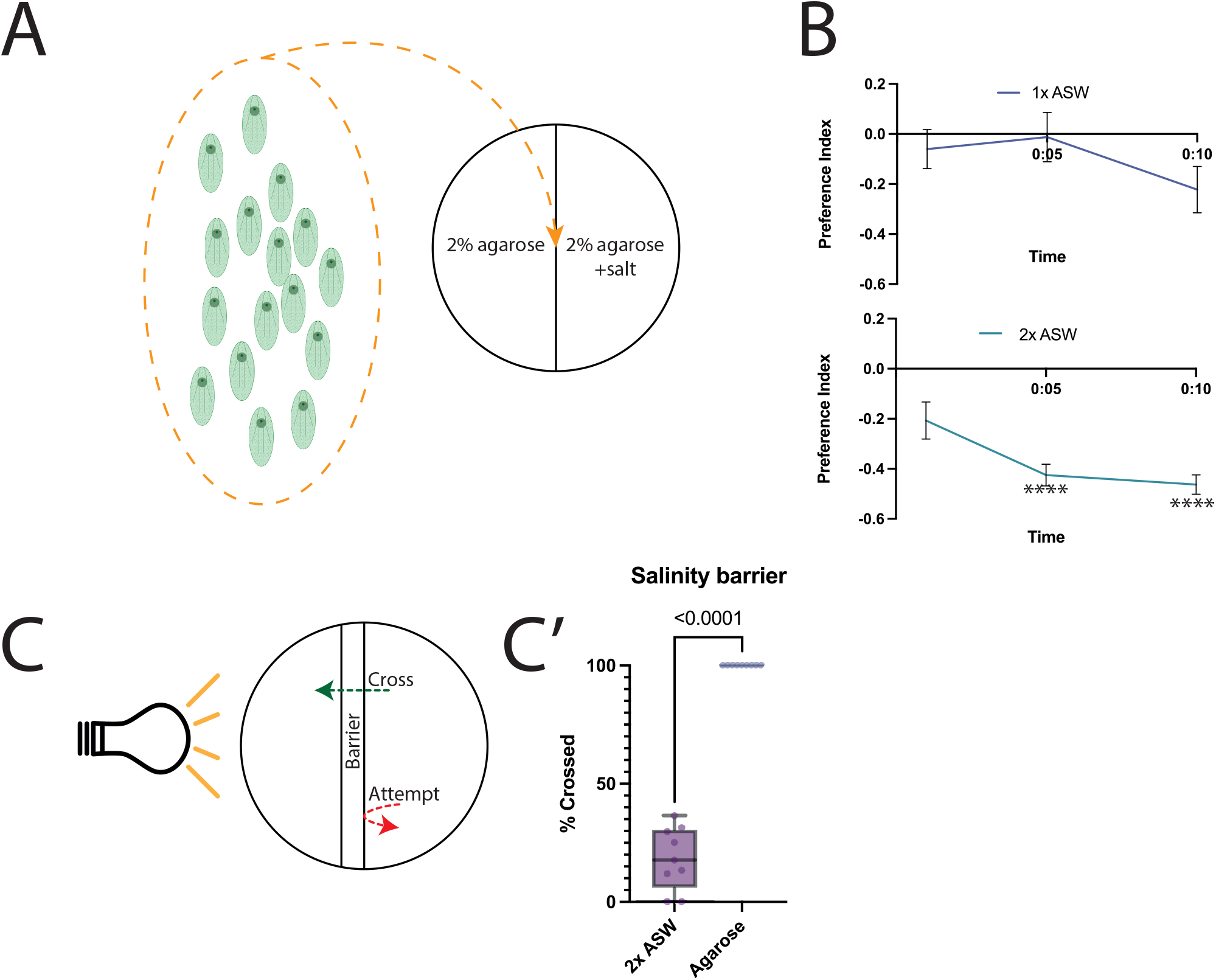
behavioural responses of *S. roscoffensis* towards varied salinity conditions. A: overview of experimental conditions showing pooled animals being placed on the behaviour arena comprised of plain agarose on one half, and agarose with added salt content on the other half. B: Time-series diagrams showing preference indices over time, with twofold ASW concentration showing significance beyond the 5-minute timepoint; N=10 for all trials, mean and SEM shown. C: salinity barrier assay diagram indicating the considered “attempt” and “cross” determinants; C’: – quantification of successful crosses across the high-salinity barrier compared with a plain agarose barrier; N=10 trials, all points and min-max bars shown. **** - p<0.0001, ns where not shown, Brown-Forsythe and Welch ANOCA test with Dinnet’s T3 multiple comparisons tested against the control group.

The finding that *S. roscoffensis* appears to avoid higher salt concentrations suggests that this condition acts as repellent cue for the animal. However, we wanted to evaluate whether the animals decided to avoid higher salt concentrations, or whether another physiological effect, such as reduced mobility, was at play. To test this, we developed a barrier-assay, in which we took advantage of the known strong positive phototaxis behavior of *S. roscoffensis*. In short, we removed a 1cm agarose strip in the middle of the plain agarose plate and replaced it with agarose containing twofold ASW. Subsequently a group of animals was placed on one side of the strip while a LED lamp was placed, laterally, at the opposing side with the aim of inducing the animals to cross the midline attracted by the light. We recorded their movement across the high-salinity barrier during phototaxis. Here, we observed that when presented with a plain (agarose in dH_2_O) barrier, 73% (403/553) of animals attempted (evaluated by counting the encounter of the animals with the barrier) to cross the barrier towards the light source, and 100% (403/403) of the attempts were successful, evaluated by the complete crossing of the barrier towards the light source. Meanwhile, when presented with a two-fold ASW barrier, 52% (184/352) of animals attempted to cross the barrier, with only 19.6% (36/184) of attempts resulting in a cross (figure 2C). Therefore, we show that despite the presence of a highly attractive light stimulus, the animals avoid the approach it if they encounter aversive conditions, suggesting a hierarchical aversion-attraction perception and decision making.

Next, we asked if adult *S. roscoffensis* are responsive to chemical cues emitted by symbiontic and non-symbiontic algae (which also live in their surrounding waters). Thus, to probe whether the presence or absence of the symbiont *T. convolutae* or other, closely related algal species (*C. concordia, T. chuii*, and *T. striata*) may be used as a sensory cue, we performed similar two-choice assays, providing the animals a choice between plain agarose and either a fresh algal culture F/2 medium, or a supernatant of an extract of algal cultures, over 10-minute trials with preference indices being measured at the 2-, 5-, and 10-minute timepoints. We observed that while there is a strong preference for pure F/2 medium, the mature animals strongly avoid the supernatants from cultures of *T. convolutae, C. concordia, T. chuii*. However, they show a slight but statistically insignificant avoidance of *T. striata* extracts (figure 3A). Furthermore, we utilised the same barrier decision-making assay to determine whether the presence of the *T. convolutae* supernatant displays similar aversive properties as compared to excessive salinity. Here, we found that only 37% (252/687) of animals attempted to cross the barrier, with a total success rate of 19.4% (49/252) (figure 3B).

**Figure 3:**
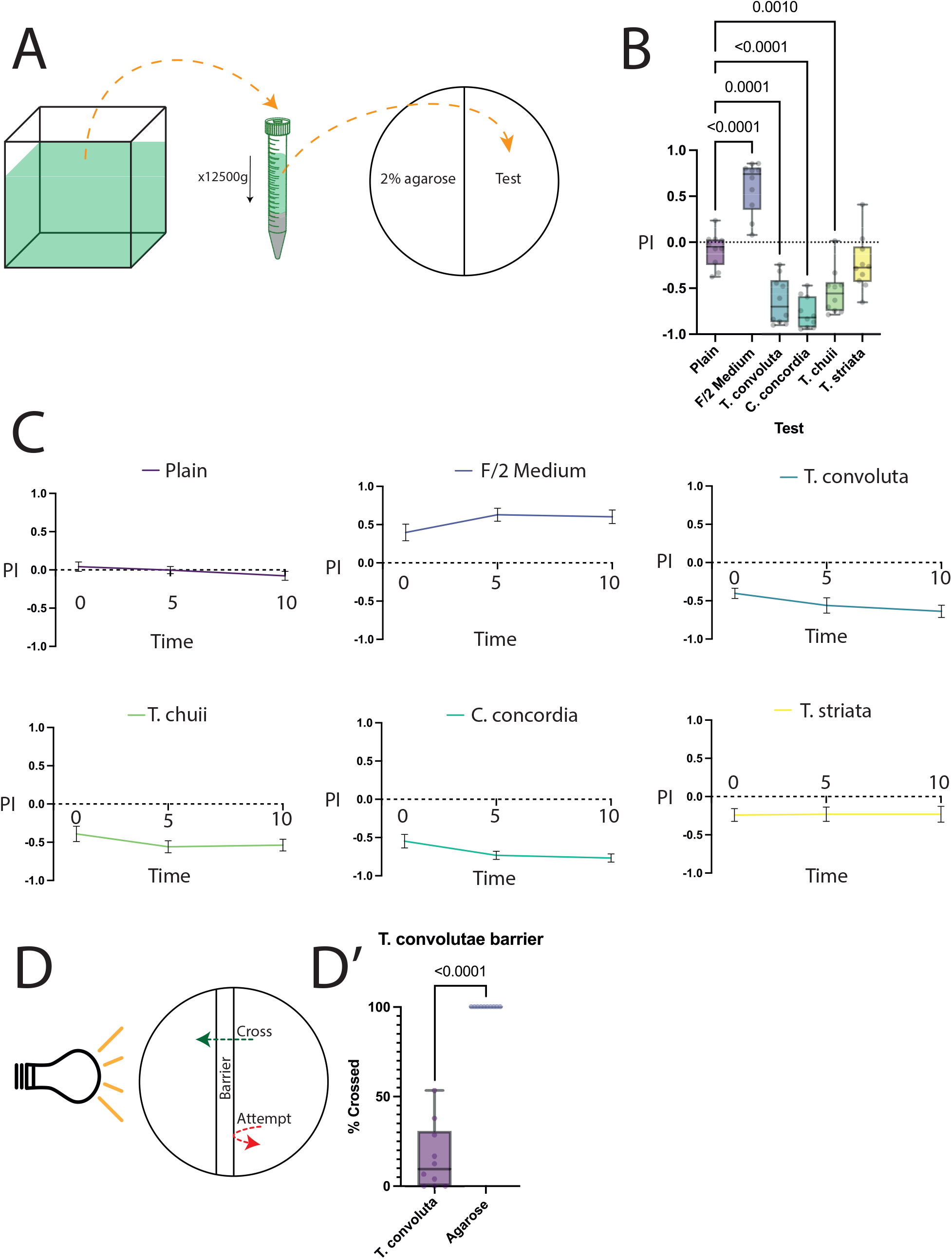
behavioural tests for algal symbiont extract. A: diagram showing the process of extraction of algal culture. Approximately 20ml of algal culture was Dounce homogenised, then spun down and supernatant was used to prepare the choice assay, with half of the plate containing plain agarose, and the other half containing agarose dissolved in extracted supernatant. B: summary of behaviour phenotypes at the 10-minute timepoint, with F/2 medium by itself showing a strong attractive index, while supernatant extracted from algal cultures shows strong aversive indices; N=10 for all trials, all points and min-max bars shown. C: time-series diagrams showing preference indices over time. F/2 medium preference is significant beyond the 5-minute timepoint, while supernatant from *T. convoliutae, T. chuii*, and *C. concordia* causes immediate strong aversion. While supernatant from *T. striata* also shows a mild aversion, it is not statistically significantly different from the plain control. N=10 for all trials, mean with SEM shown. D: *T. convolutae* culture supernatant barrier cross assay performed similarly to the salinity barrier assay. D’: *S. roscoffensis* was less likely to cross the barrier towards the light when compared with a plain agarose barrier.

Apart from chemical environmental cues, temperature may act as a key determinant for an animal to navigate towards favourable conditions. Therefore, we chose to characterise whether S. roscoffensis displays also preference for specific temperature conditions. Here, we placed groups of animals on plates with different temperature gradients: a) high (30^°^C) to low (5^°^C), b) high to ambient (18^°^C), c) low to ambient, or d) ambient to ambient ranges (figure 4A). We observed that there is, as expected, no difference in the side preference between ambient-to-ambient areas (10min, PI=-0.01), and, similarly, no significant preference between the low to ambient conditions (10min, PI=-0.05, P=0.9366 vs ambient control). On the other hand, when presented with a choice between a high and low temperature, the animals tend to prefer the low temperature side (10min, PI=-0.37, P=0.0028 vs ambient control). Moreover, the animals preferred the side with ambient temperature in comparison to the side with higher temperature (10min, PI=-0.30, P=0.0398 vs ambient control), suggesting an avoidance behaviour towards higher temperatures (figure 4B, C). These experiments support the notion that the animals do prefer lower temperatures over ambient or high temperatures. This in turn suggests that while low temperatures are neither attractive nor aversive, high temperature is decidedly an aversive cue.

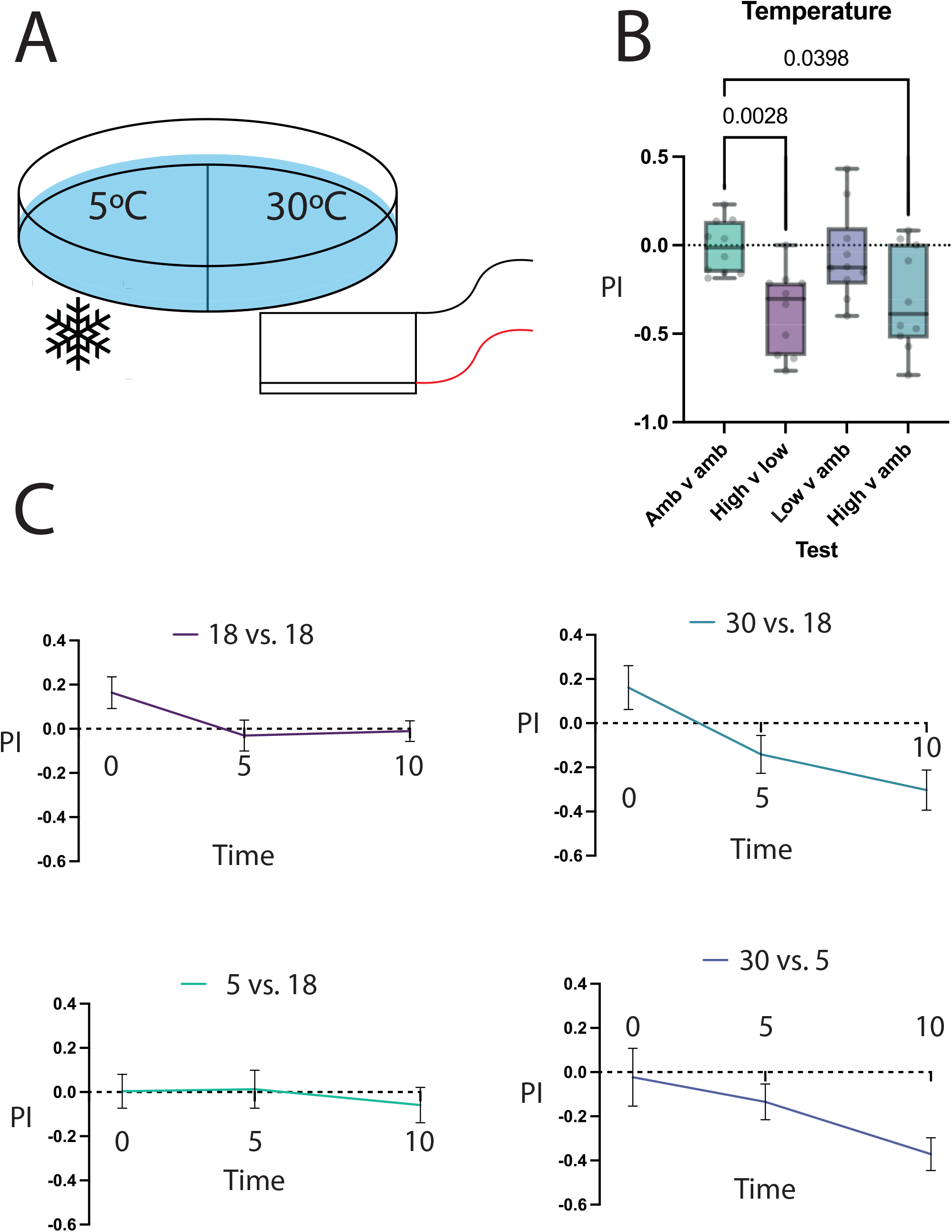

## Discussion

Intertidal animals are exposed to variable environmental conditions, whether daily or along the seasons (Adey and Loveland, 2007). These conditions have generated specific adaptations, reflected in their physiologies and their behavioural responses. Here we analysed the preferences to chemical and physical cues of the acoel flatworm *Symsagittifera roscoffensis*, an acoel living in obligatory symbiosis most of their life. Our findings indicate that the acoel *Symsagittifera roscoffensis* is able to display distinct behavioural preferences when exposed to a range of chemical and environmental stimuli. Furthermore, the variability of chemicals used in our assays allows us the uncovering of varied set of phenomenologies. For instance, the differing responses to different amino acids raise a question about the mechanistic nature of this response. One explanation of the positive preference towards Tryptophan, a precursor to serotonin and melatonin, may be due to either a direct behavioural decision to seek it out due to regular scarcity, or alternatively, it may be due to a continuous need in the bioavailability of serotonin in the brain, where it could modulate (as in other invertebrates) basic aspects of their physiology such as feeding, olfactory sensitivity, egg laying, and homeostasis.

The aversion to increased salinity may also relate to the naturally varying changes in the habitat of the worms. During low tide, as water evaporates from the surface of the sand, salinity will increase, thus causing a changing environment for the animals. Thus, it is logical that the animals would prefer a specific level of salinity to remain in an isotonic environment and avoid a hypotonic one. However, it is known that *S. roscoffensis* emerges to the surface during low tide conditions, thus contraindicating an aversion to increasing salinity levels. One explanation for this may be that the strong negative geotaxis, but not positive phototaxis responses, overpower the natural aversion to increased salt levels, and the worms instead fine-tune their positioning within different levels of salinity on the surface.

The results for the responses to the symbiont supernatant also provide a surprising new perspective on the interactions between these host (worm) and the algae. While it is known that juvenile *S. roscoffensis* incorporate algae into their bodies, suggesting a level of attraction to any chemical signals that the algae may emit, adults show the opposite response. Firstly, we controlled for whether it is the medium of the algal culture that causes a behavioural phenotype, and, indeed, we found that it is an appetitive stimulus. However, despite this, the presence of any algal emissions, and not the algae themselves, causes a strongly aversive response. It is possible to speculate that once the animals reach a state of maturity, including incorporating enough algae within them (with a narrow stoichiometry), a physiological switch occurs which causes them to avoid the free-living alga in their immediate environment. Perhaps, this may be in order to avoid overcrowding and allow new or growing juveniles more access to the algae, thus ensuring that the population is maintained, and self-competition is avoided. On the other hand, considering that in these experiments algal extract was used, the aversive behaviour may be due to the animals avoiding broken (thus dead) algae, perhaps being expelled by dead animals. The presence of certain chemicals may thus indicate that the locality is under stress, as indicated by the liberated chemicals and metabolites from broken algae, indicating that the animals need to hide or migrate to safer places (e.g. DMSP, dimethylsulphoniopropionate (Van Bergeijk and Stal, 2001)). Further studies on adult and juvenile *S. roscoffensis* may provide the answers to these questions.

Like the responses of the salinity assays, the phenotype of high-temperature avoidance may also play a role in the navigation of the animals on the surface during low tide. Since increasing temperature causes increased evaporation of the water, this may be an interlinked response with the avoidance of high salinity. Furthermore, higher temperatures may result in increased metabolism within the algae, or the worms themselves. Increased metabolism may result in increased stress, for example from excess oxygen production resulting in elevated oxidative stress. Thus, avoiding increased temperatures may be a physiological response to avoid excess or premature death. Contrarily, lower temperatures may instead result in a neutral preference index due to either a lack of ability to sense such temperatures, or the fact that the temperature of the water fluctuates between an average low of 9^°^C to an average high of 17^°^C across the year. Thus, the “low temperature” condition in our assay may not be sufficiently lower than the temperature experienced by *S. roscoffensis* in the natural environment. In all, we show that it is possible to pose a question that the interplay between a variety of stimuli combinations result in the natural behaviour of *S. roscoffensis*, opening an avenue for further investigation.

## Material and Methods

### Animal collection and maintenance

*S. roscoffensis* was collected from wild colonies from the sands in the vicinity of Roscoff and Carantec (Brittany, France). The collected sand was then placed in deep trays and *S. roscoffensis* was collected with Pasteur pipettes during low tides when the animals display negative geotaxis. Collected animals were transferred into plastic containers with natural sea water, continuously flowing, at the Station Biologique de Roscoff (Roscoff, Brittany, France). All experiments were performed during similar tidal and circadian conditions, between 09:00 and 15:00.

### Two-choice behaviour assays

All behaviour assays were performed using the method described in Maier et al., 2021, substituting *S. roscoffensis* for *Drosophila* larvae. In brief, 87mm petri dishes were filled with 2% agarose diluted in distilled water and allowed to solidify for a minimum of 10 minutes. Following this, one half of the agarose layer was cut and discarded. Then, the now-empty half of the dish was filled with 2% agarose containing the substance of choice diluted in distilled water (for details, see below). *S. roscoffensis* were pooled in groups between 50-200 individuals and placed in the centre of the plate, with enough sea water being added to form a complete, thin, layer on top of the agarose. Experiments were monitored and the number of animals in each half-plate were counted after 2, 5 and 10 minutes. The animals on each side were then counted and a preference index was calculated using the following formula:

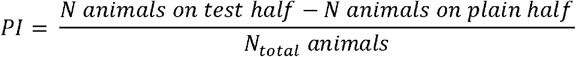

The chemicals used were of the highest purity available, as follows:

Amino Acid assays: L-Glutamine (Sigma-Aldrich, 56-85-9), L-Aspartic acid (Sigma-Aldrich, 56-84-8), L-Tyrosine (Sigma-Aldrich, 60-17-4), L-Tryptophan (Sigma-Aldrich, 73-22-3), L-Glycine (Roth, 3187.3). All tests were performed at 20mM concentration.

Artificial sea water (Aquarium Sytems, Instant Ocean) diluted in distilled water to a concentration of 34ppm (onefold) or 68ppm (twofold).

Choice assays with culture supernatant from the classical and facultative symbiont: *Tetraselmis convolutae* (the classical symbiotic algae of *S. roscoffensis*), and *Tetraselmis striata, Tetraselmis chuii* and *Chlamydomonas concordia*, (three facultative symbionts of *S. roscoffensis*) were cultured in flasks containing of sea water and F/2 feeding medium (UTEX culture collection of algae) for a period of 15 days. Before collecting the supernatant, the algae were examined at the microscope to exclude any abnormalities or increased cell death. The content of the flasks was collected by means of a Douce tissue grinder following 20 passes of the pestle, before centrifuging the slurry. 87mm petri dishes were prepared with one half containing 2% agarose in distilled water and the other half containing 2% agarose in a 25 ml mix composed of 12 ml distilled water and 13 ml supernatant solution of one of each four algae species, respectively.

### Barrier assays with twofold salinity and culture supernatant from the classical symbiont

87mm dishes were prepared with 2% agarose in distilled water on both sides, with a 10mm strip cut out from the middle, and subsequently filled with agarose containing twofold ASW or *T. convolutae* supernatant. An LED lamp (OSARAM LED, 80012 White) was placed at a 45° angle, 9 cm above the table at a distance such that the lamp-oriented side of the dish was illuminated at 1100 lux, the middle of the plate 840 lux and the side opposite of the lamp with 400 lux (Keene et al., 2011; von Essen et al., 2011). Animals were placed on the side of the barrier further from the light, and subsequently observed over the course of 10 minutes, with the attempts and successful crosses counted.

### Thermal preference assays

87mm petri dishes were coated with 2% agarose before being placed in a Styrofoam box for thermal insulation. One half of the petri dish was in contact either with ice water (reaching a temperature of 5^°^C), RT (18-21^°^C) or with a Peltier plate running at 24W, reaching a temperature of 30^°^C. Animals were pooled and placed in the middle of the petri dish in a similar fashion to the choice assays described above. The number of animals was then recorded on each side, and a preference index was calculated.

## Acknowledgements

We would like to express our gratitude to the Swiss National Science Foundation (SNSF) for providing the funding for this project (grant 310030_219348 and IZKSZ3_218514 to SGS). We are also thankful to the University of Fribourg (UniFr), members of the Department of Biology and the Marine Station in Roscoff for their support and help throughout our research. We also thank all members of the Sprecher Lab for their invaluable feedback on our experimental design, procedures, and reporting.

## References

Achatz, J.G., Chiodin, M., Salvenmoser, W., Tyler, S., Martinez, P., 2013. The Acoela: on their kind and kinships, especially with nemertodermatids and xenoturbellids (Bilateria incertae sedis). Org. Divers. Evol. 13, 267–286. 10.1007/s13127-012-0112-4

Adey, W.H., Loveland, K., 2007. CHAPTER 2 - The Envelope: Physical Parameters and Energy State, in: Adey, W.H., Loveland, K. (Eds.), Dynamic Aquaria (Third Edition). Academic Press, London, pp. 13–42. 10.1016/B978-0-12-370641-6.50011-X

Arboleda, E., Hartenstein, V., Martinez, P., Reichert, H., Sen, S., Sprecher, S., Bailly, X., 2018. An Emerging System to Study Photosymbiosis, Brain Regeneration, Chronobiology, and Behavior: The Marine Acoel Symsagittifera roscoffensis. BioEssays 40, 1800107. 10.1002/bies.201800107

Bailly, X., Laguerre, L., Correc, G., Dupont, S., Kurth, T., Pfannkuchen, A., Entzeroth, R., Probert, I., Vinogradov, S., Lechauve, C., Garet-Delmas, M.-J., Reichert, H., Hartenstein, V., 2014. The chimerical and multifaceted marine acoel Symsagittifera roscoffensis: from photosymbiosis to brain regeneration. Front. Microbiol. 5.

Bailly, X., Martinez, P., Hartenstein, V., Gavilán, B., Simon G. S., 2021. Symsagittifera roscoffensis as a Model in Biology, in: Handbook of Marine Model Organisms in Experimental Biology. CRC Press.

Bery, A., Cardona, A., Martinez, P., Hartenstein, V., 2010. Structure of the central nervous system of a juvenile acoel, Symsagittifera roscoffensis. Dev. Genes Evol. 220, 61–76. 10.1007/s00427-010-0328-2

Boyle, J.E., Smith, D.C., 1975. Biochemical Interactions between the Symbionts of Convoluta roscoffensis. Proc. R. Soc. Lond. B Biol. Sci. 189, 121–135.

Dingledine, R., McBain, C.J., 1999. Glutamate and Aspartate Are the Major Excitatory Transmitters in the Brain, in: Basic Neurochemistry: Molecular, Cellular and Medical Aspects. 6th Edition. Lippincott-Raven.

Geddes, P., 1879. II. Observations on the physiology and histology of Convoluta schultzii. Proc. R. Soc. Lond. 449–457. 10.1098/rspl.1878.0153

Geng, X., Boufadel, M.C., Jackson, N.L., 2016. Evidence of salt accumulation in beach intertidal zone due to evaporation. Sci. Rep. 6, 31486. 10.1038/srep31486

Keeble, F., 1911. Plant-Animals: a Study in Symbiosis. Nature 86, 446–446. 10.1038/086446b0

Lanfranchi, A., 1990. Ultrastructure of the epidermal eyespots of an acoel platyhelminth. Tissue Cell 22, 541–546. 10.1016/0040-8166(90)90082-K

Maier, G.L., Komarov, N., Meyenhofer, F., Kwon, J.Y., Sprecher, S.G., 2021. Taste sensing and sugar detection mechanisms in Drosophila larval primary taste center. eLife 10, e67844. 10.7554/eLife.67844

Martínez, P., Hartenstein, V., Sprecher, S.G., 2017. Xenacoelomorpha Nervous Systems, in: Oxford Research Encyclopedia of Neuroscience. 10.1093/acrefore/9780190264086.013.203

Philippe, H., Brinkmann, H., Copley, R.R., Moroz, L.L., Nakano, H., Poustka, A.J., Wallberg, A., Peterson, K.J., Telford, M.J., 2011. Acoelomorph flatworms are deuterostomes related to Xenoturbella. Nature 470, 255–258. 10.1038/nature09676

Roshchina, V.V., 2016. New Trends and Perspectives in the Evolution of Neurotransmitters in Microbial, Plant, and Animal Cells, in: Microbial Endocrinology: Interkingdom Signaling in Infectious Disease and Health. Springer, Cham, pp. 25–77. 10.1007/978-3-319-20215-0_2

Sakagami, T., Watanabe, K., Hamada, M., Sakamoto, T., Hatabu, T., Ando, M., 2024. Structure of putative epidermal sensory receptors in an acoel flatworm, Praesagittifera naikaiensis. Cell Tissue Res. 395, 299–311. 10.1007/s00441-024-03865-y

Selosse, M.-A., 2000. Un exemple de symbiose algue-invertébré à Belle-Isle-en-Mer: la planaire Convoluta roscoffensis et la prasinophycée Tetraselmis convolutae. Acta Bot. Gallica 147, 323–331. 10.1080/12538078.2000.10515864

Sprecher, S.G., Bernardo-Garcia, F.J., van Giesen, L., Hartenstein, V., Reichert, H., Neves, R., Bailly, X., Martinez, P., Brauchle, M., 2015. Functional brain regeneration in the acoel worm Symsagittifera roscoffensis. Biol. Open 4, 1688–1695. 10.1242/bio.014266

Thomas, N.J., Tang, K.W., Coates, C.J., 2024. To move or not to move: taxis responses of the marine acoel Symsagittifera roscoffensis to different stimuli. Mar. Freshw. Behav. Physiol. 57, 17–31. 10.1080/10236244.2024.2337444

Todt, C., Tyler, S., 2007. Ciliary receptors associated with the mouth and pharynx of Acoela (Acoelomorpha): a comparative ultrastructural study. Acta Zool. 88, 41–58. 10.1111/j.1463-6395.2007.00246.x

Van Bergeijk, S.A., Stal, L.J., 2001. Dimethylsulfoniopropionate and dimethylsulfide in the marine flatworm Convoluta roscoffensis and its algal symbiont. Mar. Biol. 138, 209–216. 10.1007/s002270000444

Yamasu, T., 1991. Fine structure and function of ocelli and sagittocysts of acoel flatworms. Hydrobiologia 227, 273–282. 10.1007/BF00027612

